# Quantitative Assessment of 3D Printed Blood Vessels Produced with J750^™^ Digital Anatomy^™^ for Suture Simulation

**DOI:** 10.1101/2022.01.09.475308

**Authors:** Stefania Marconi, Valeria Mauri, Erika Negrello, Luigi Pugliese, Andrea Pietrabissa, Ferdinando Auricchio

**Affiliations:** Department of Civil Engineering and Architecture, University of Pavia, Pavia, Italy; Foundation IRCCS Policlinico San Matteo, Pavia, Italy; Department of Clinical, Surgical, Diagnostic and Paediatric Sciences, University of Pavia, Pavia, Italy

## Abstract

Blood vessels anastomosis is one of the most challenging and delicate tasks to learn in many surgical specialties, especially for vascular and abdominal surgeons. Such a critical skill implies a learning curve that goes beyond technical execution. The surgeon needs to gain proficiency in adapting gestures and the amount of force expressed according to the type of tissue he/she is dealing with. In this context, surgical simulation is gaining a pivotal role in the training of surgeons, but currently available simulators can provide only standard or simplified anatomies, without the chance of presenting specific pathological conditions and rare cases.

3D printing technology, allowing the manufacturing of extremely complex geometries, find a perfect application in the production of realistic replica of patient-specific anatomies. According to available technologies and materials, morphological aspects can be easily handled, while the reproduction of tissues mechanical properties still poses major problems, especially when dealing with soft tissues.

The present work focuses on blood vessels, with the aim of identifying – by means of both qualitative and quantitative tests - materials combinations able to best mimic the behavior of the biological tissue during anastomoses, by means of J750^™^ Digital Anatomy^™^ technology and commercial photopolymers from Stratasys. Puncture tests and stitch traction tests are used to quantify the performance of the various formulations. Surgical simulations involving anastomoses are performed on selected clinical cases by surgeons to validate the results.

A total of 37 experimental materials were tested and 2 formulations were identified as the most promising solutions to be used for anastomoses simulation. Clinical applicative tests, specifically selected to challenge the new materials, raised additional issues on the performance of the materials to be considered for future developments.

## 1. The role of 3D Printing in surgical training

Surgical simulation is gaining a pivotal role in the training of surgeons [1–4], in order to guarantee a balance between patient’s safety and a proper acquisition of technical skills, self-awareness and self-confidence by young surgeons. Available simulators – based on the use of animals, cadavers, or standard mannequin – can provide only standard or simplified anatomies, without the chance of presenting specific pathological conditions and rare cases, thus preventing the training on the possible anatomical-related challenges of the clinical practice.

3D printing (3DP) technology is gaining popularity in many fields, such as automotive, aerospace, architecture and fashion; however, the medical and dental sectors [5] are growing extremely fast, thanks to the chance of personalization and customization offered by 3DP technologies. 3DP allows the manufacturing of extremely complex geometries, thus perfectly suitable for producing realistic replica of patient-specific anatomies. While morphological aspects can be easily handled, the **reproduction of tissues mechanical properties poses major problems**, especially when dealing with soft tissues, such as blood vessels.

**Blood vessels anastomosis** is one of the most challenging and delicate tasks to learn in many surgical specialties, especially for vascular and abdominal surgeons. Such a critical skill implies a learning curve that goes beyond technical execution. The surgeon needs to gain proficiency in adapting gestures and the amount of force expressed according to the type of tissue he/she is dealing with, in terms of thickness and elasticity of the blood vessels wall, presence of calcific plaques etc. Today advances in mini-invasive surgery are leading to an increasing spread of laparoscopic and robotic instruments, which are becoming an almost mandatory part of novice surgeons’ background. In this context, the absence of a tactile feedback can make the task even more complex to accomplish. The surgeon in this case needs to rely on the visual feedback, thus tuning the force applied to the tissue according to its visual deformation. Therefore, it appears clear how the material used for producing anatomical replicas intended for training must be carefully studied to **grant a realistic behavior**.

In this context, photopolymers represent the most promising class of 3DP materials. The availability of highly deformable photopolymers combined with Material Jetting technology - which makes it possible to locally mix materials – enables a fine tuning of the final object’s mechanical properties. The high resolution of Material Jetting technology also grants the reproduction of the finest morphological details.

The present work focused on the use of the **J750^™^ Digital Anatomy^™^**, one of the latest innovative releases by Stratasys®. This Material Jetting printer, more specifically a PolyJet 3D printer, has a wide selection of available materials, ranging from opaque or transparent materials to flexible or biocompatible photopolymer resins, which can be mixed to a number of combinations. It features a minimum layer thickness of 14 microns and a build size of 1400 x 1260 x 1100 mm.

Being oriented to the medical field applications, J750^™^ Digital Anatomy^™^ enables the use of three unique digital materials and an extensive library of *anatomical presets*, that is predefined combinations of materials studied to create a range of biomechanical models that look, feel, and respond like real biological tissues.

In particular, innovative dedicated resins are:

- Bone Matrix: thanks to the deposition of specific patterns, it is able to mimick porous bone structures, fibrotic tissues or ligaments which provide realistic feedback when cutting and drilling;
- Gel Matrix: it allows to print the smallest, most complex and thin vascular structures and easily remove internal support material;
- Tissue Matrix: the softest 3D printing material available on the market which allows to produce organs’ models that mechanically behave like the real anatomical structures.

Thanks to these features, the J750^™^ Digital Anatomy^™^ is the best candidate for the development of innovative solutions for the surgical simulation field.

**In the literature, mechanical properties of 3DP materials are only assessed through qualitative studies, based on the subjective feedback of surgeons** on the dissection, cutting and suturing of 3D printed anatomical models. Although various examples of experimental set-ups for the quantitative evaluation of mechanical parameters are available in the literature - such as resistance to suturing or to surgical needles penetration [6–19] - these studies always concern biological (human and animal) or synthetic materials, while no works evaluate the performance of 3DP materials.

**The present work focuses on blood vessels, with the aim of identifying – by means of both qualitative and quantitative tests - materials combinations able to best mimic the behavior of the biological tissue during anastomoses**. More in detail, main goals are (i) the development of a methodology for a quantitative evaluation of the performance of 3DP materials obtained combining various photopolymer resins; (ii) the set-up of specific mechanical tests to assess 3DP materials performance in mimicking vessels’ mechanical response during suturing; (ii) the identification of the 3DP materials able to best mimic the various vessels’ response during suture, by means of comparison with biological samples.

## 2. Methodology

### 2.1 From qualitative to quantitative assessment of mechanical performances

An expert surgeon – with more than 15 years of experience in abdominal surgery – was initially enrolled to identify the main qualitative parameters of interest and the best surgical thread to perform sutures. The choice fell on a non-absorbable polypropylene/polyethylene 5/0 monofilament with a 3/8 18mm cylindrical needle (PROLENE^®^ - Ethicon), commonly used to suture small-to-medium caliper abdominal vessels (in the following Prolene 5/0). The choice was made in order to test the selected 3DP materials in one of the worse use cases: the thinner the filament, the more the 3DP material will be prone to tearing under suturing. Rectangular samples (100×20×1-2 mm) made of different photopolymer combinations were produced through a J750^™^ Digital Anatomy^™^. All the samples were printed using *Matte finishing* and *Heavy Support* options, since these are the recommended print conditions for the production of patient-specific anatomical models. After printing, samples were manually cleaned to remove most of the support structure and then placed for 2 hours in the cleaning solution (2% sodium hydroxide and 1% sodium metasilicate (Na_2_SiO_3_)). Afterwards, they were rinsed under tap water and then placed in a 15% technical-grade glycerol (99% purity level) solution for 30 seconds. Samples were left drying before testing for at least 12 hours at room temperature.

For each material, the surgeon performed a continuous suture - with distinct points at first stitch - using the selected Prolene 5/0 surgical thread on each sample cut in halves. The filament has a diameter of about 0.13 mm. Distance between stitches usually depends on the thickness of the specific vessel. For major blood vessels, with a thickness above 1 mm and an inner diameter of 10-20 mm, distant stitches are used, while for smaller vessels (below 1 mm of thickness and below 10 mm of inner diameter) closer stitches are usually required.

According to surgeon’s comments during the suturing exercises, the following parameters were identified as valuable for 3D printed samples performance evaluation: (i) puncture force, (ii) resistance to the stitch, (iii) thread sliding, (iv) surgical knots tightening and (v) juxtaposition of the segments.

From the mechanical point of view, puncture force and resistance to stitch were considered the most relevant and therefore analyzed also from a quantitative point of view. An experimental *ad hoc* set-up was designed to translate qualitative parameters (i) and (ii) into quantitative mechanical measures.

**Puncture tests** were performed to measure the force required to pierce the material with the surgical needle. On the other hand, uniaxial traction tests were performed on sutured samples and on samples with a single stitch applied for **resistance to stitch** evaluation. Single stitch test was selected to avoid any influence coming from the surgeons’ peculiar approach during suturing. Details of each test will be discussed in the next paragraphs.

All the mechanical tests were performed through an MTS Insight Testing System^®^ with a 250N load cell. The development of the various components of the experimental set-up was based both on the available scientific literature [6–19] and on the ASTM F1342/F1324M-05 (2013) standard related to the resistance to penetration of materials for protective clothing [20]. All the components were designed on Autodesk Inventor^®^ CAD software and 3D printed. The same mechanical tests were performed also on porcine aorta samples, to be used as a biological comparison.

### 2.2 Puncture Tests

For the measure of the puncture force, cylindrical samples – 40 mm diameter and thickness ranging from 1 to 2 mm - were produced and tested through the *ad hoc* experimental set-up designed to measure puncture force (Figure 1).

**Figure 1:**
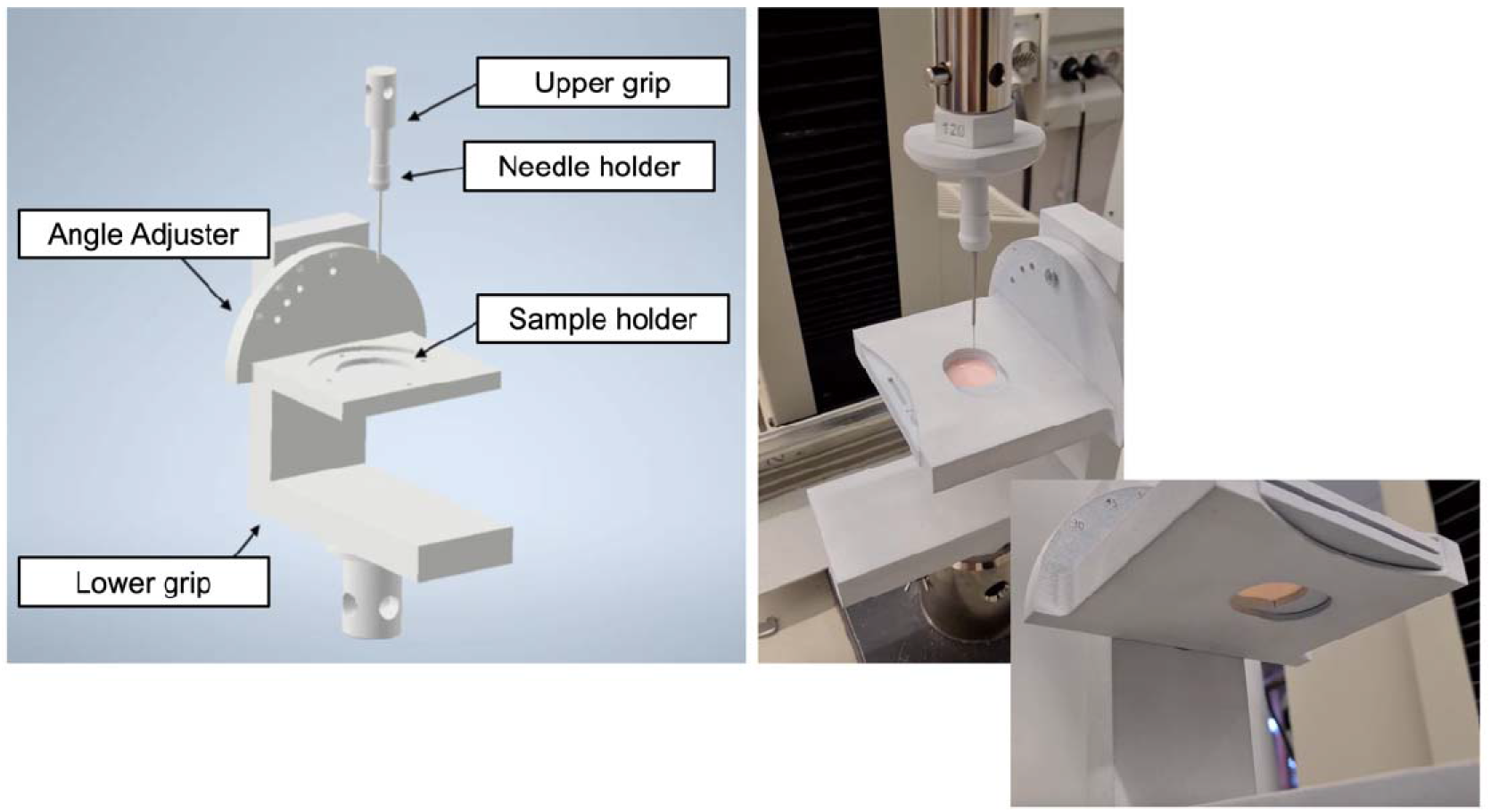
Experimental set-up for puncture mechanical tests.

To correctly perform the test with the MTS Insight, it was necessary to use a rectilinear needle; after an unsuccessfully manual rectification on the needle of the selected Prolene 5/0 surgical thread, a rectilinear needle with the most similar characteristic to Prolene 5/0 was selected (Trusilk USP 4/0, a rectified 19mm cylindrical needle).

In the literature, most of the studies analyzing materials’ puncture resistance use common punches or needles to perform mechanical tests. On the contrary, in the present study only specific surgical needles were employed to consider the actual dynamic of laceration these needles have on biological tissues. In fact – unlike common needles or punches – surgical needles are purposely made to be as fine as possible to minimize trauma and to be sharp enough to penetrate tissue with minimal resistance.

Some vascular anastomosis videos were analyzed to estimate the average speed of the needle passing the tissue during stitches; by slowing down the videos, dividing it into different frames and considering the thickness of the sutured vessels, a medium speed of 5mm/s was estimated. To carry out the puncture tests, a specific set up to mount the samples and the surgical needle on the MTS Insight Testing System^®^ was designed and produced. A mechanical compression tests with an acquisition rate of 20Hz, a crosshead speed of 5mm/s and a 250N load cell was employed to characterize the performance of the different 3D printed materials.

### 2.3 Single Stitch & Suture Tests

Both for single stitch and suture tests Prolene 5/0 was used as well. In the first case, a single stitch is made approximately at 1,7 mm from the margin. The actual distance was then measured with a caliper and recorded. For suture tests, two surgeons – with more than 15 and 5 years of experience respectively - performed the suture on the different 3D printed samples. Rectangular samples (100×20×1-2 mm) were used in both tests. Each sample was cut in halves and then a single stitch or multiple stitches were applied according to the test.

All the samples were subjected to uniaxial tensile test with an acquisition rate of 20Hz, a crosshead speed of 5mm/s and a 250N load cell.

**Figure 2:**
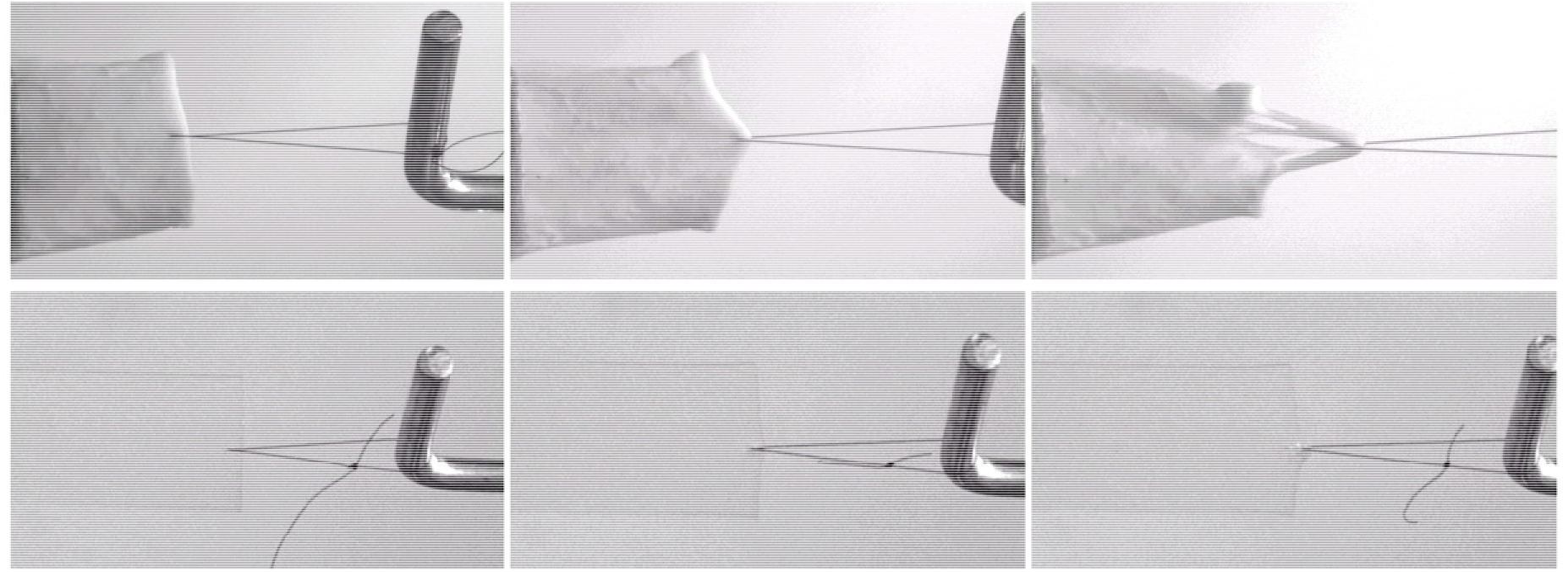
Comparison between single stitch mechanical tests performed on porcine aorta (up) and on a 3DP material (down).

**Figure 3:**
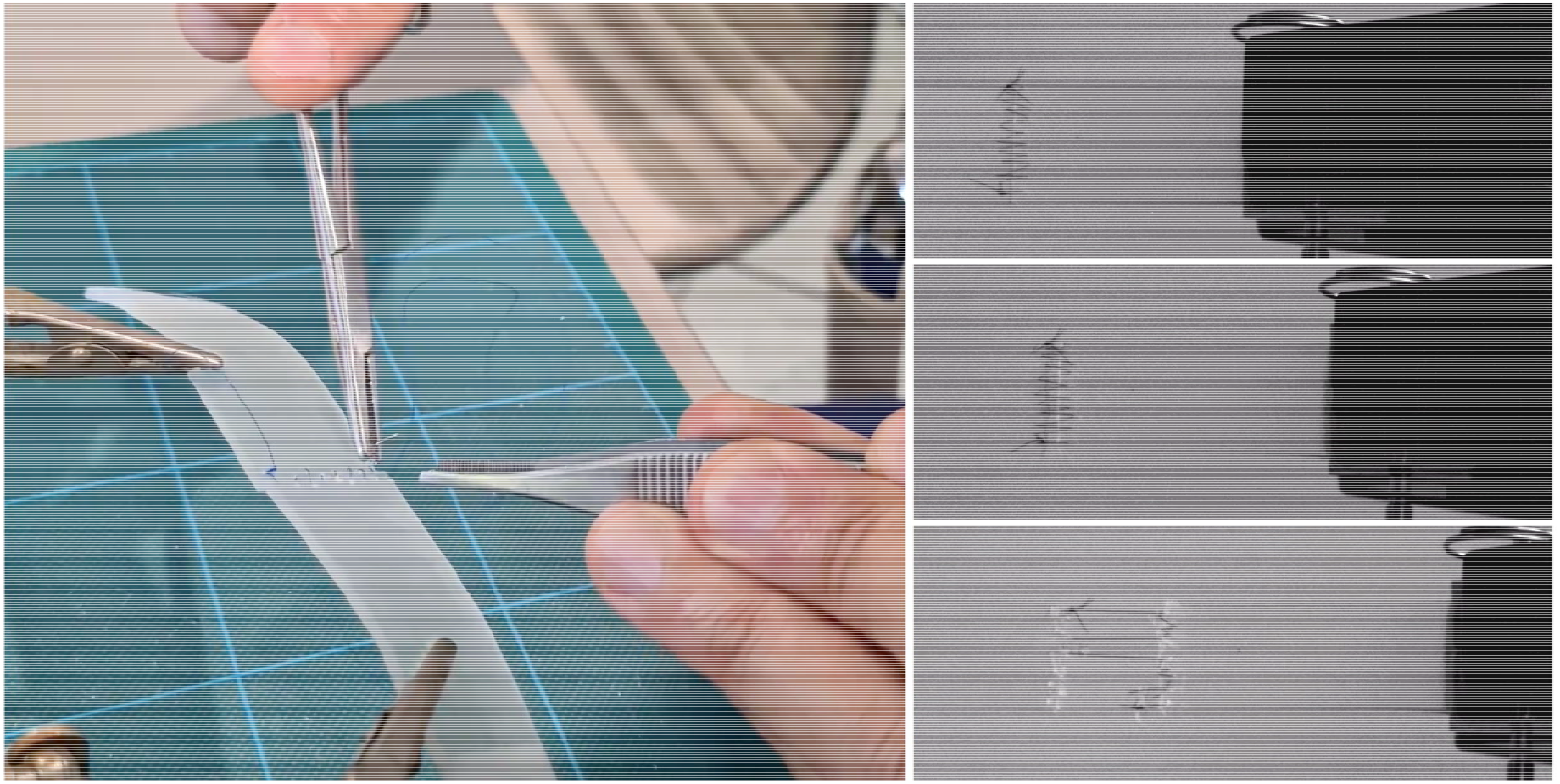
Uniaxial tensile test performed on a 3D printed sample sutured by an expert surgeon in order to mimic the surgical suture on a medium caliper arterial vessel.

### 2.4 Benchmark with biological samples

Two fresh porcine aortas were tested and used as a biological comparison for the tested 3DP materials.

Both aortas were cleaned with pliers and surgical scalpels within 2 hours from the explant, in order to remove the tissue surrounding the vessels and were kept in 0,9% saline solution at 37° throughout the cleaning and testing time.

From each aorta various samples were obtained, divided according to the aortic tract; in particular, for all the aortas, 4 samples of aortic arch, 6 samples of thoracic aorta, 6 samples of abdominal aorta, 4 samples for lower abdominal aorta and 2 for renal arteries were obtained.

**Figure 4:**
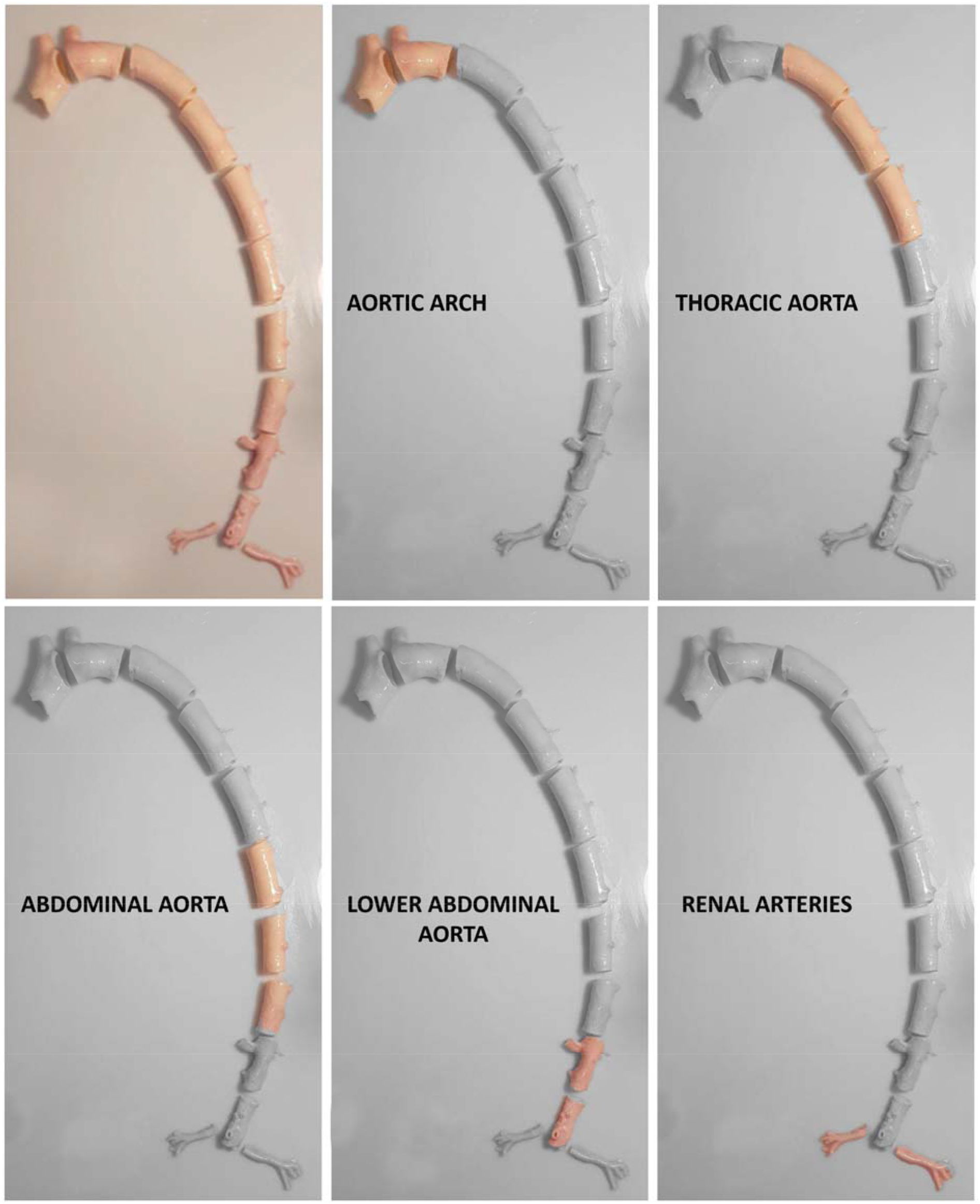
Preparation of porcine aorta samples of the various vascular tracts.

Biological specimens were subject to the **puncture** and **single stitch** tests previously described, with the same experimental set-up, instrumentation and testing parameters used to test the various 3DP materials. Puncture test is performed only on aortic arch, thoracic and abdominal aorta, while single-stitch test is performed also on the lower abdominal tract and renal arteries. The thickness of each sample is recorded after the test in the immediate proximity of the region of the test (i.e. near the puncture and near the single stitch)

## 3. Results

A total of 37 different material–combinations have been tested for both puncture and single stitch uniaxial traction tests. A final pool of 6 material combinations was thoroughly tested with 10 repetitions performed for each material and mechanical test type, for a total of more than 120 mechanical tests performed on the final set of materials, both for puncture and traction tests. Moreover, uniaxial traction tests were also performed on sutured samples.

In the following, the pool of 6 3D printing materials will be referred to as A, B, C, D, E, F.

Among these 6 materials, according to mechanical tests results and surgeons’ feedbacks, 3 materials were considered as the best performing ones. The results of the previous described mechanical characterization tests are presented below. First, an overview of the results on the 6 selected materials is presented (A-F). Then more details are given for the 3 final materials.

### 3.1 Overall Results

Results from puncture and single-stitch tests are here presented in terms of the following parameters (Figure 5).

- Ultimate Stress 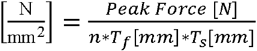 The highest displacement acquired during the single-stitch test divided by the surface of the filament in contact with the material. Given the low thickness of the filament used – 0,13 mm – the surface was simplified into a rectangular area computed as the thickness of the filament (*T_f_*) multiplied by thickness of the sample (*T_s_*);
- Normalized Max Displacement 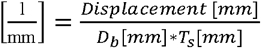 The highest displacement acquired during the single-stitch test divided by the product of the distance of the stitch from the border (*D_b_*) and the thickness of the sample (*T_s_*);
- Unit Peak Force 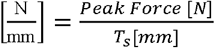 The highest force acquired during the puncture test divided by the thickness of the sample.

**Figure 5:**
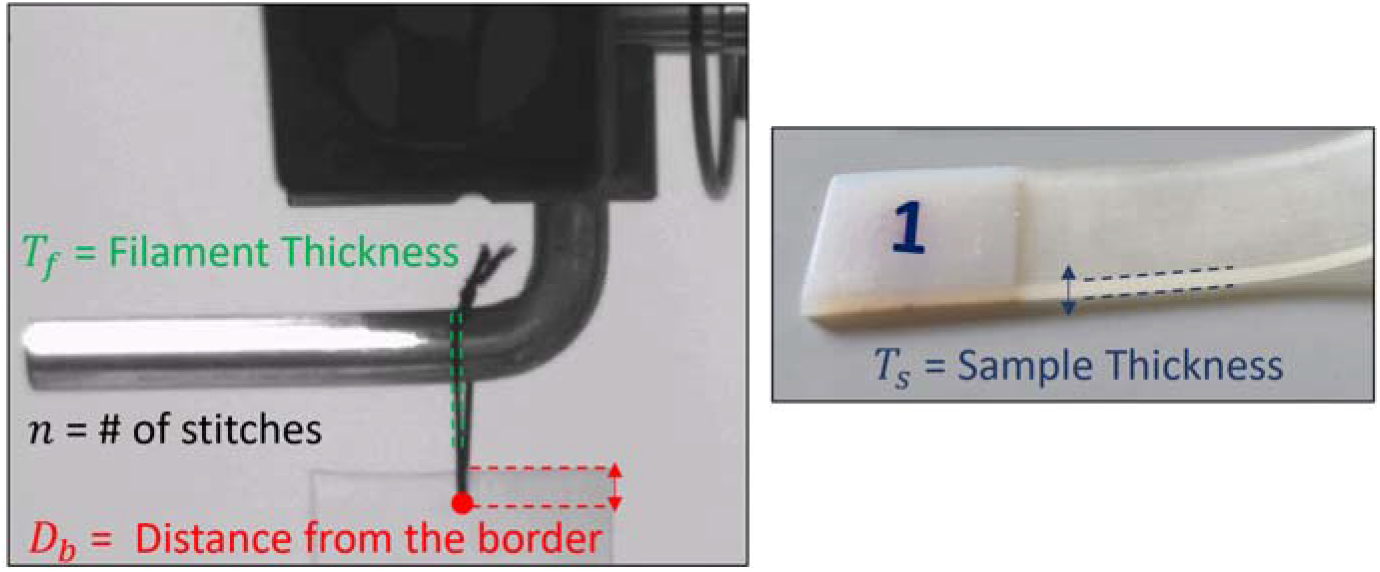
Scheme of the parameters employed in the normalization of the tests results.

These parameters are computed with the aim of making the result of each test independent from the features of the specific sample.

In Figure 6 all the data collected on porcine samples during single stitch and puncture tests are presented. It is clearly visible how the results on biological tissues are affected by data dispersion: this is due to the continuous local change in the biological tissue structure moving from the aortic arch to the abdominal aorta, and to the presence of a variable amount of residual biological tissue on the vessels’ outer wall. Being puncture tests available only for aortic arch, thoracic aorta and abdominal aorta, and being lower abdominal tract and renal arteries extremely different in terms of performance under single-stitch traction tests (see Figure 9), the comparison was limited to the first three segments of the aorta.

**Figure 6:**
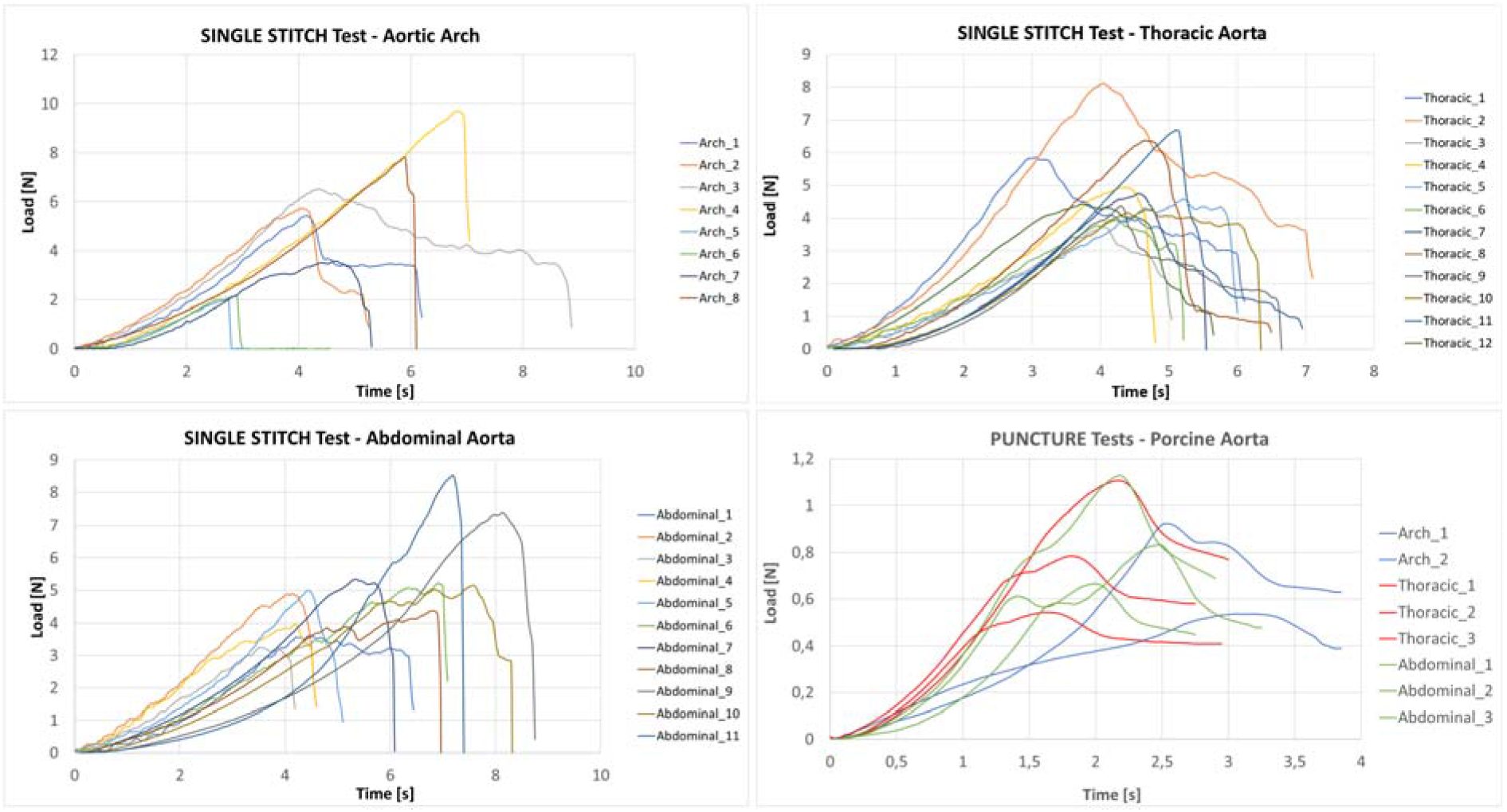
Results of single stitch and puncture tests on all the porcine aorta samples.

**Figure 7:**
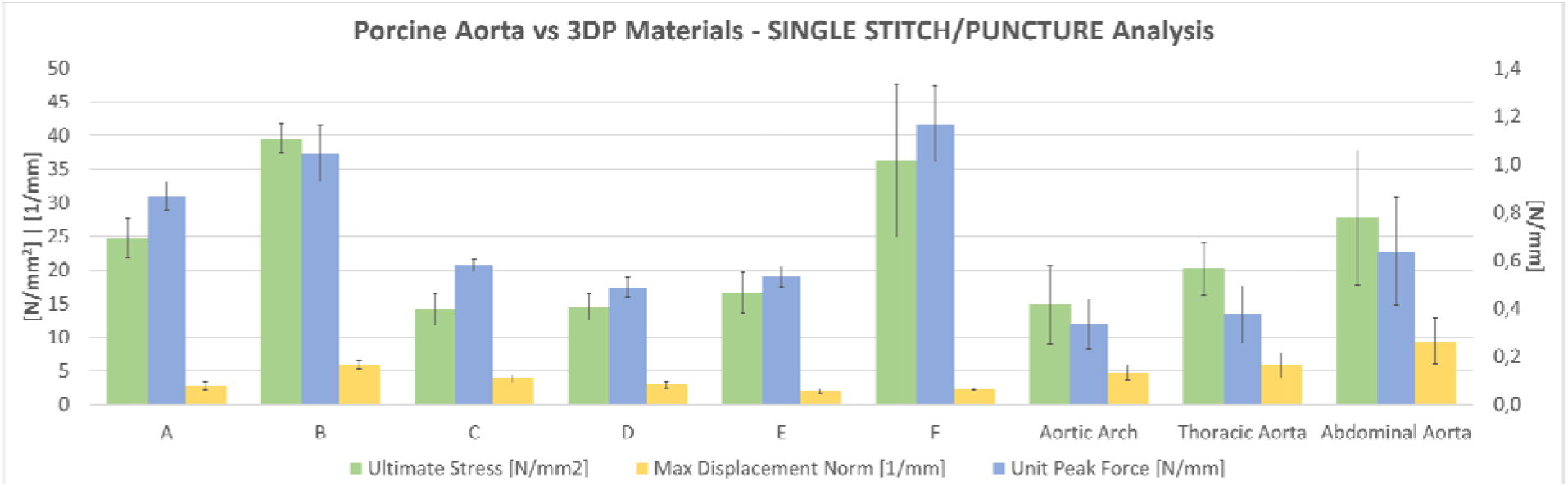
Overall results of puncture and single-stitch tests on the pool of 6 selected materials compared with porcine aorta tracts (namely aortic arch, thoracic aorta and abdominal aorta).

**Figure 8:**
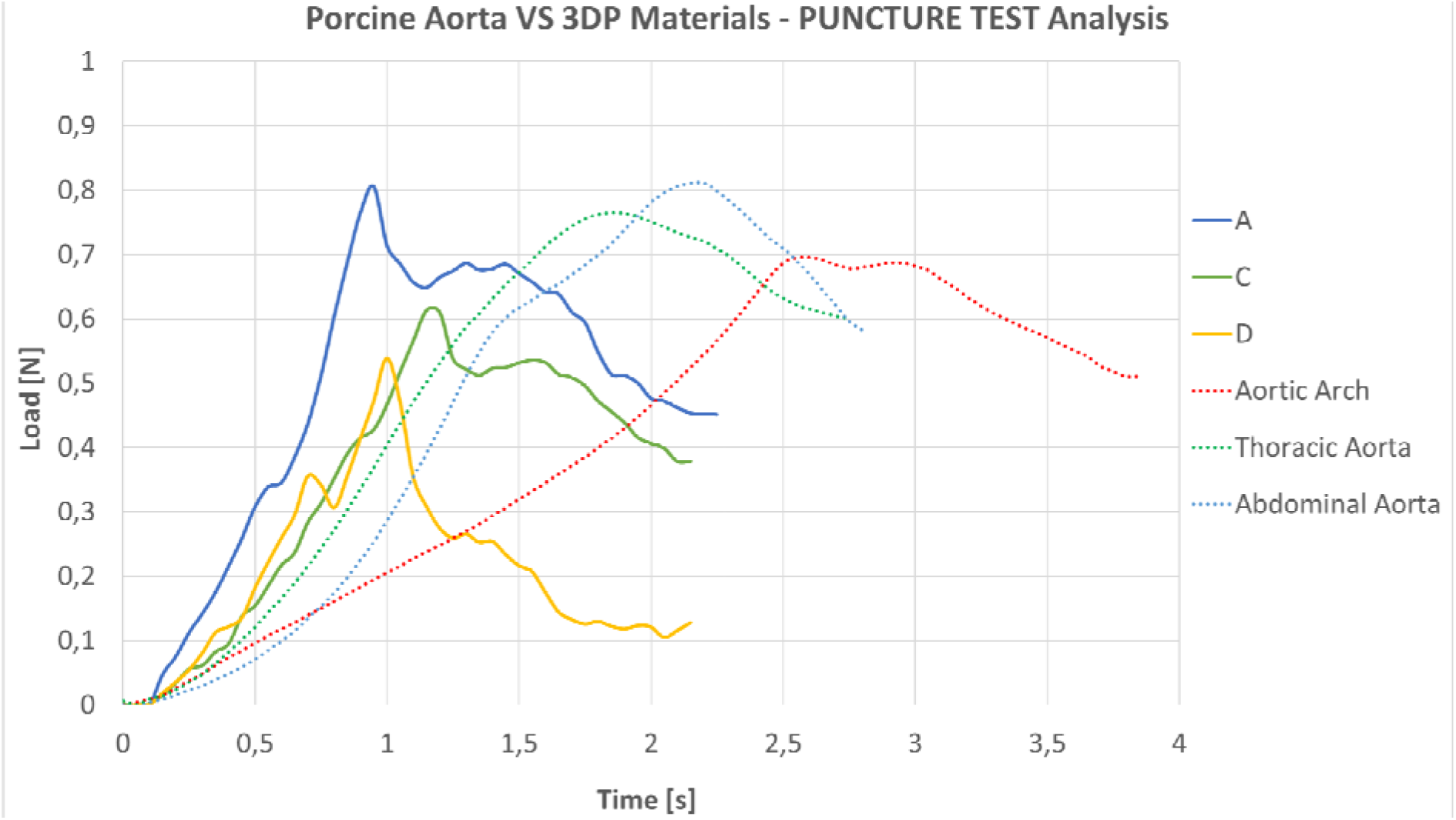
Load-time curves acquired during puncture tests: comparison between 3DP materials and different porcine aorta tracts on one representative curve per material.

**Figure 9:**
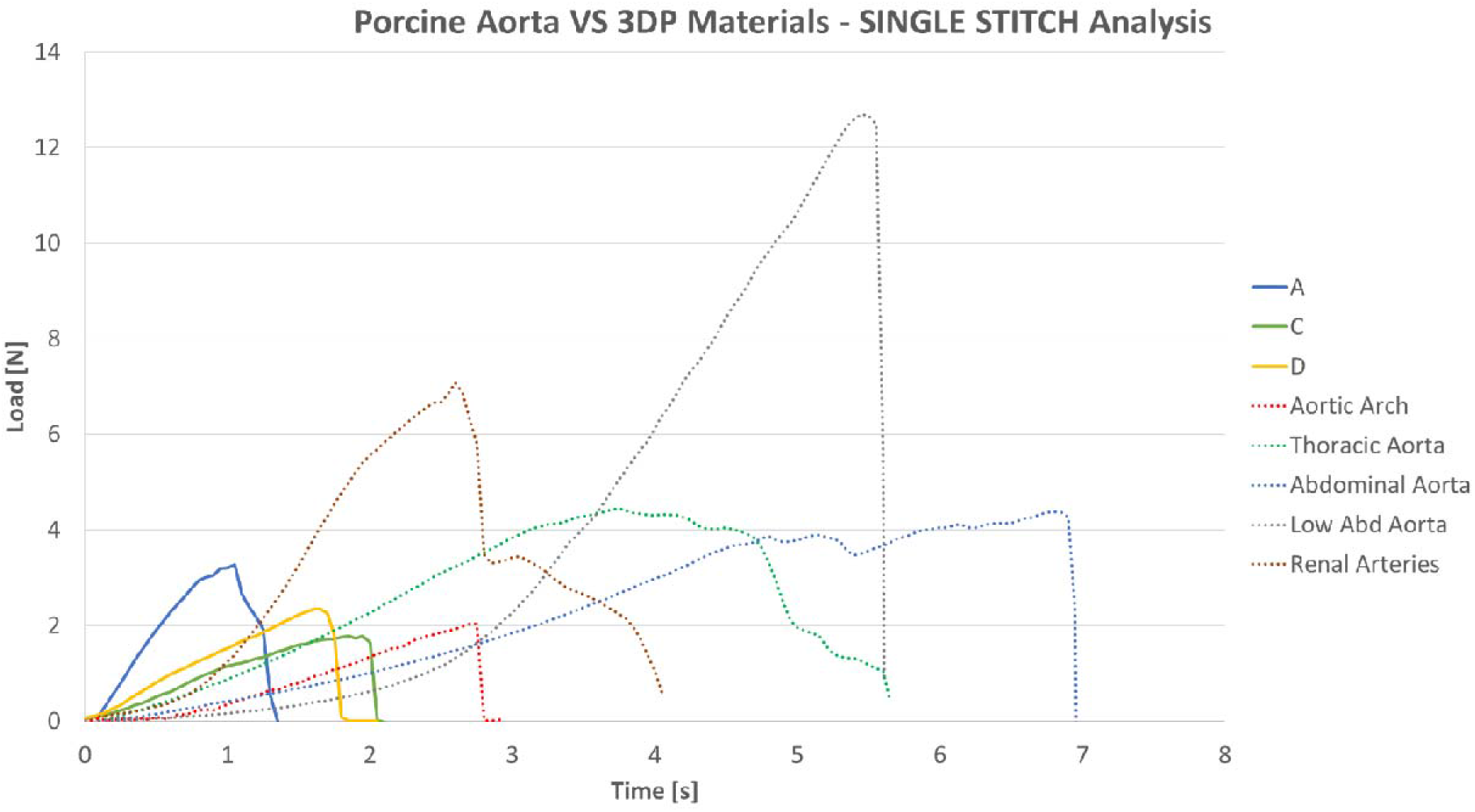
Load-time curves acquired during single stitch mechanical tests: comparison between 3DP materials and different porcine aorta tracts on one representative curve per material.

Figure 7 shows the overall results on the final pool of selected materials, compared to the porcine aorta According to the results, B and F were excluded for the extremely high values in terms of puncture force. As it is visible in Figure 2, the displacement of the biological tissue is way higher than the average displacement of 3D printed materials. On the other hand, this parameter plays a fundamental role in the replica of the feedback that the tissue gives to the surgeon during suturing. The other 4 materials were also subjected to a qualitative evaluation with the surgeon, to confirm their performance.

A, C, D and E underwent the qualitative suture test performed by the 2 expert surgeons, as described in Section 2.

Both surgeons rated the 4 materials as comparable in terms of puncture force. Unit Peak Force values range between 0,49 and 0,87 N/mm. E was rated as the worst in term of friction of the filament during sliding through the material, and the worst in terms knots tightening and resistance to stitch, since visible tearing around the stitches was present. The juxtaposition of the segments was discarded during the last tests since it was considered too dependent from the specific set-up, which can possibly force unrealistic positioning of the two segments. No specific drawbacks were highlighted during the tests.

**According to surgeons’ feedback, A, C and D were selected as the most interesting materials for suture simulation**. In the following, more detailed information about the mechanical performances of these 3 materials are presented.

### 3.2 Puncture Tests

Puncture tests performed on A, C and D show mean Unit Peak Force of 0.87 ± 0.06 N/mm, 0.58 ± 0.03 N/mm and 0.49 ± 0.04 N/mm respectively. On the other hand, same tests on the different aortic tracts provided values of 0.34 ± 0.1 N/mm for the aortic arch, 0.38 ± 0.12 N/mm for the thoracic aorta and 0.64 ± 0.22 N/mm for the abdominal aorta.

Figure 8 reports the load-time curves, in which values are not normalized according to the sample thickness; for clarity, normalized values are reported in the table below.

**Table 1:** Normalized puncture tests’ results: comparison between 3DP materials and different porcine aorta tracts.

| Materials | Thickness [mm] | #Samples | Puncture Tests |
| --- | --- | --- | --- |
|  |  |  | Unit Peak Force [N/mm] |
| **A** | **1** | **5** | **0,87 ± 0,06** |
| **C** | **1** | **4** | **0,58 ± 0,03** |
| **D** | **1** | **5** | **0,49 ± 0,04** |
| AORTIC ARCH | 2,2 ± 0,26 | 2 | 0,34 ± 0,10 |
| THORACIC AORTA | 2,18 ± 0,19 | 3 | 0,38 ± 0,12 |
| ABDOMINAL AORTA | 1,46 ± 0,19 | 3 | 0,64 ± 0,22 |

Porcine aorta samples have higher thickness with respect to the 3DP samples, which are 1mm thick, thus Unit Peak Force values should be considered for numerical comparison. Table 1 highlights that normalized numerical values of the 3DP materials and biological samples unit peak force are comparable, while Figure 8 shows how the peak values of biological samples occur later than in 3D printed materials: this result is in line with the single-stitch behavior visible in Figure 2, due to the higher elasticity of the biological tissue, which can hold much higher elongation before rupture.

### 3.3 Single Stitch Tests

Single stitch mechanical tests performed on A, C and D show mean ultimate stress of 24.64 ± 2.92 N/mm^2^, 14.15 ± 2.35 N/mm^2^ and 14.48 ± 2.03 N/mm^2^ respectively. The results of mean ultimate stress for the different biological samples are: 14.82 ± 5.85 N/mm^2^ for the aortic arch, 20.17 ± 3.84 N/mm^2^ for the thoracic aorta, 27.74 ± 10.06 N/mm^2^ for the abdominal aorta, 12.7 ± 0.72N for the lower abdominal aorta and 10.02 ± 4.17 for the renal arteries.

Figure 9 reports the load-time curves, in which values are not normalized according to the sample thickness. For greater clarity and for a more reliable comparison – based on different samples’ thicknesses -, normalized values are reported in the table below.

**Table 2:** Normalized single stitch tests’ results: comparison between 3DP materials and different porcine aorta tracts.

| Materials | Thickness [mm] | # Samples | Single Stitch Traction Tests |  |  |  |
| --- | --- | --- | --- | --- | --- | --- |
|  |  |  | Peak Force [N] | Ultimate Stress [N/mm<sup>2</sup>] | Max Displacement [mm] | Max Displacement Normalized [1/mm] |
| A | 1 | 10 | $3,20 \pm 0,38$ | $24,64 \pm 2,92$ | $6,10 \pm 0,92$ | $2,77 \pm 0,60$ |
| C | 1 | 10 | $1,84 \pm 0,31$ | $14,15 \pm 2,35$ | $8,10 \pm 1,32$ | $3,85 \pm 0,57$ |
| D | 1 | 10 | $1,88 \pm 0,26$ | $14,48 \pm 2,03$ | $6,40 \pm 1,17$ | $2,94 \pm 0,49$ |
| AORTIC ARCH | $2,2 \pm 0,26$ | 8 | $5,40 \pm 2,32$ | $14,82 \pm 5,85$ | $24,40 \pm 8,02$ | $4,71 \pm 1,19$ |
| THORACIC AORTA | $2,18 \pm 0,19$ | 12 | $5,20 \pm 1,35$ | $20,17 \pm 3,84$ | $22,44 \pm 3,43$ | $5,87 \pm 1,82$ |
| ABDOMINAL AORTA | $1,46 \pm 0,19$ | 11 | $5,27 \pm 1,28$ | $27,74 \pm 10,06$ | $29,48 \pm 5,87$ | $9,42 \pm 3,48$ |

While peak load of 3DP materials could be comparable to the one of the aortic arch, the main differences occurs for what concerns the samples’ elongation; while A, C and D have a normalized maximum displacement of 2.77 ± 0.6 mm^-1^, 3.85 ± 0.57 mm^-1^ and 2.94 ± 0.49 mm^-1^, the displacement for the various porcine aorta samples can reach 9.42 ± 3.48 mm^-1^ for the abdominal aorta or 12.61 ± 3.39 mm^-1^ for the renal arteries.

### 3.4 Surgeon Suture Tests

Surgeon suture tests performed on 3DP materials show a mean Ultimate Stress of 19.11 ± 3.73 N/mm^2^ for A, 10.45 ± 1.97 N/mm^2^ for C and 7.25 ± 0.58 N/mm^2^ for D, with normalized maximum elongation of 8.21 ± 2.37 mm^-1^, 23.72 ± 5.78 mm^-1^ and 11.73 ± 0.57 mm^-1^ respectively. Data have been normalized on the base of the medium distance of the stitches from the samples’ border and on the number of stitches the two surgeons used to complete the suture, as described in Section 3.1.

Figure 10 shows the good repeatability of the test, although each surgeon used a different number of stitches made at different distance from the border.

**Figure 10:**
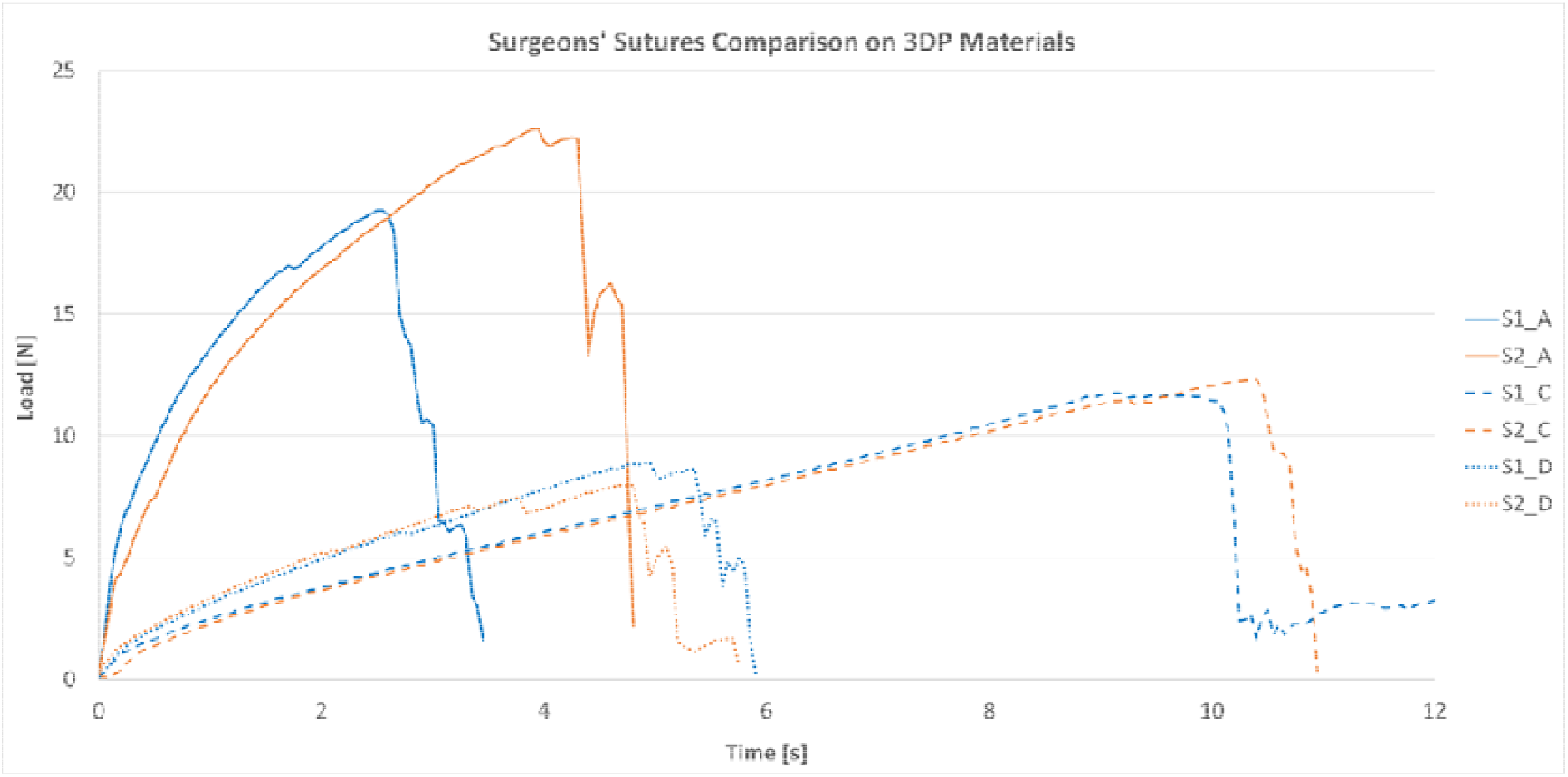
Results of surgeon sutures’ mechanical characterization: comparison between 2 surgeons (S1 and S2) with different years of experience, who completed the suture on the various 3DP materials. Each surgeon used a different number of stitches made at different distance from the border.

### 3.5 Production Tests

After mechanical tests, the 3 selected materials underwent to some applicative testing, to identify possible problems during the production of more complex structures. Patient-specific geometries used for the tests will be described in the next Section. Tests were carried out using all the different support strategies available for vascular structures: Lite/Standard/Heavy Support and GelSupport (only for the inner lumen).

- **D was excluded during these tests**, since it was discovered to be too fragile to withstand the cleaning from the support, regardless to the strength and type of the selected support structure, the vessel caliper (tested from 1-2 mm to more than 50 mm) and the wall thickness (tested up to 2,5-3 mm). Moreover, it was found to be impaired by the use of GelSupport, in vessels with variable inner diameter (tested from 1-2 mm to more than 50 mm).
- **C** was always successfully cleaned using Lite/Standard/Heavy Support in low caliper vessels (<10 mm), while it underwent to small ruptures during the cleaning of major caliper vessels (> 25 mm) of thickness below 2 mm. It was instead successfully finished with variable using GelSupport for inner lumen. For this material, the use of GelSupport is highly recommended: its used was tested also on vessels of very high caliper (aneurismatic sac of >50 mm diameter) without any impairment of the printing.
- **A** was successfully cleaned with all the support strategies and all types of vessels. GelSupport was tested also on vessels of very high caliper (aneurismatic sac of >80 mm diameter) without any impairment of the printing.

## 4. Discussion

In order to assess the performance of the selected materials for surgical simulation, 3 clinical applicative cases were selected, involving abdominal and vascular surgeons. All the models were reconstructed from Computed Tomography (CT) images and they concern real cases for which a 3DP model was asked for surgical planning purposes to 3D4Med (www.3d4med.eu), the Clinical 3D Printing Laboratory of San Matteo Hospital and University of Pavia (Pavia, Italy).

### 4.1 Aortic arch aneurism open surgery

The clinical case presents an aortic arch aneurism with a maximum diameter of 54 mm. The simulated intervention consists in the opening of the aneurismatic sac, the placement of a synthetic graft (Dacron Silver) and the closure of the aneurismatic sac over the graft (Figure 12). The anatomical model is produced with a vessel wall of variable thickness as shown in Figure 11. Two models were tested using A and C respectively: calcific plaques were printed using the “Blood Vessel – Vascular Calcification – Stiff Calcium Deposits” preset available in GrabCAD Print software for J750^™^ Digital Anatomy^™^. The model was successfully printed with GelSupport in the inner lumen, despite the high lumen diameter. The stitches were made with a Prolene 4/0 surgical thread. Moreover, the surgical repair of the left subclavian artery was tested with a synthetic patch (Figure 13). Procedures were carried out by two vascular surgeons.

**Figure 11:**
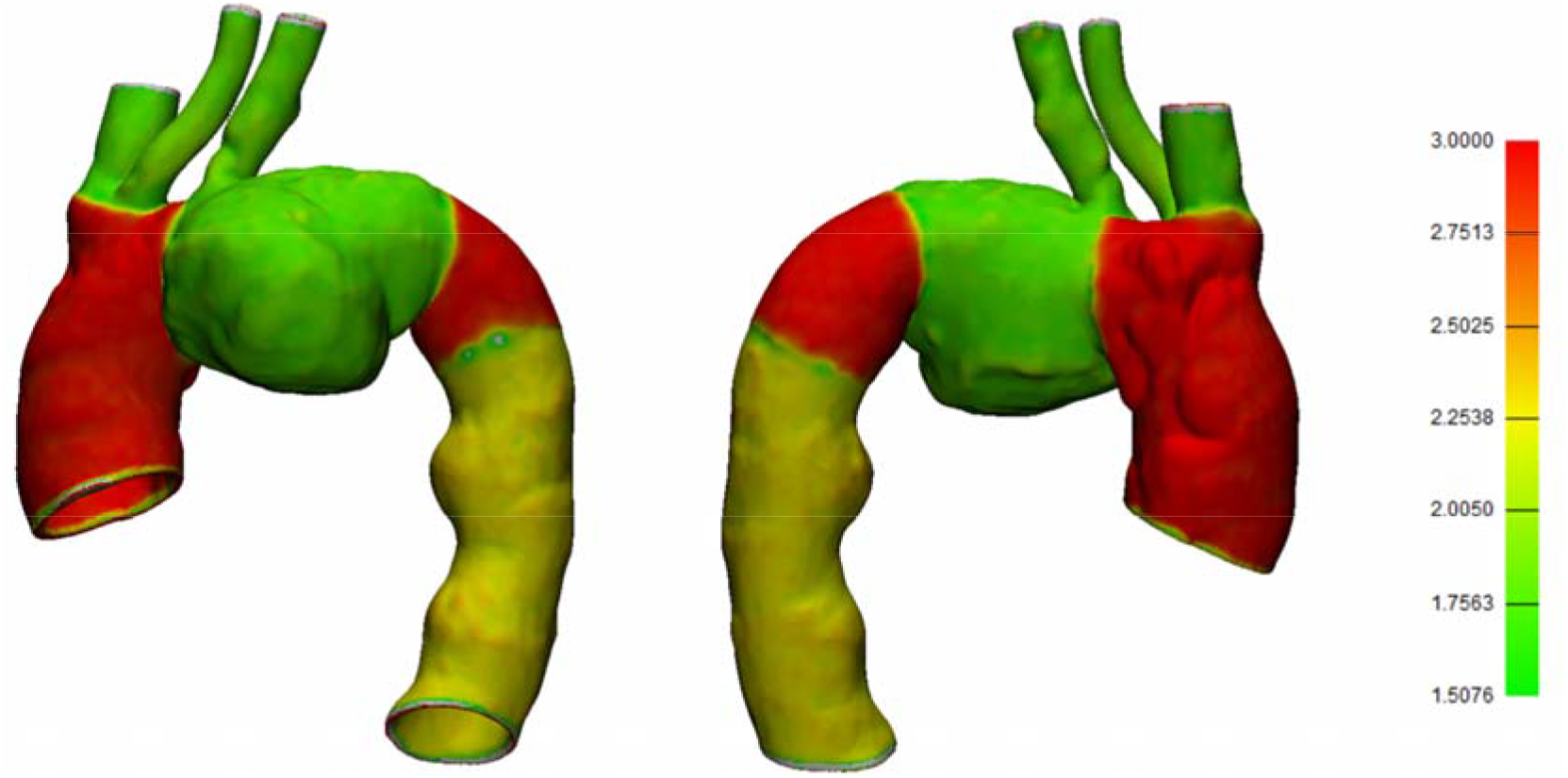
Aortic arch aneurism model wall thicknesses.

**Figure 12:**
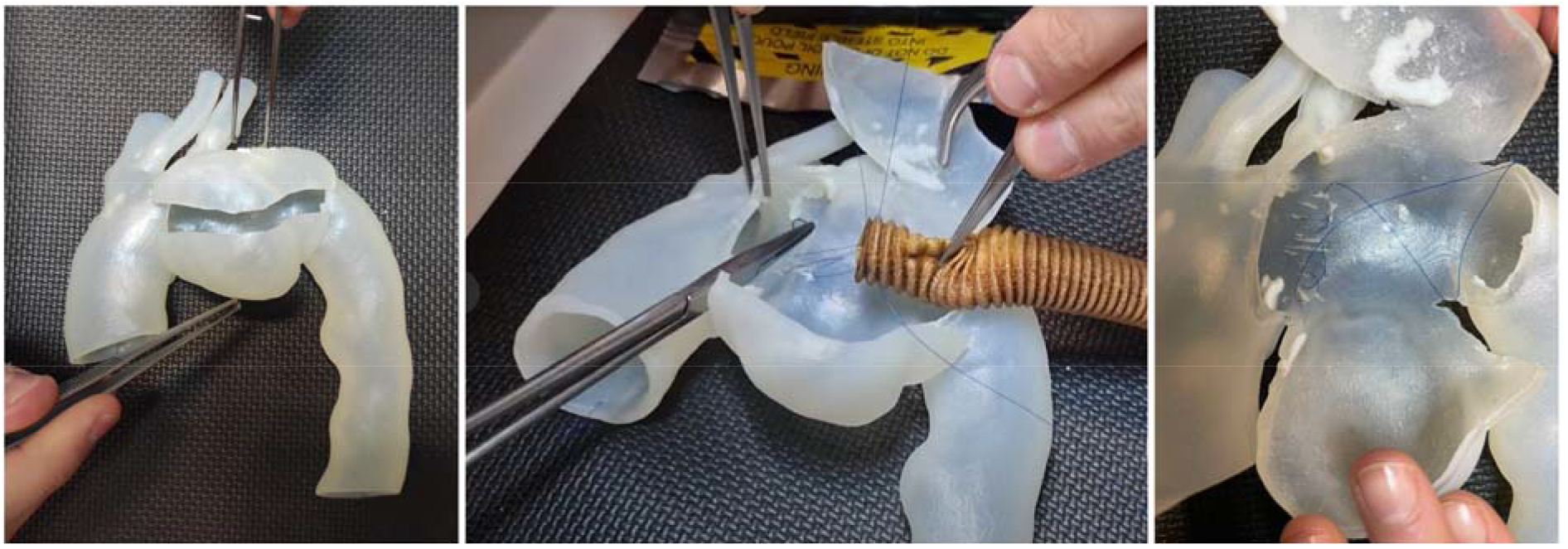
Two steps of the simulated procedure on A: opening of the aneurismatic sac (left) and placement of a synthetic graft (center). On the right, the tearing around the stitches is visible (white lines).

**Figure 13:**
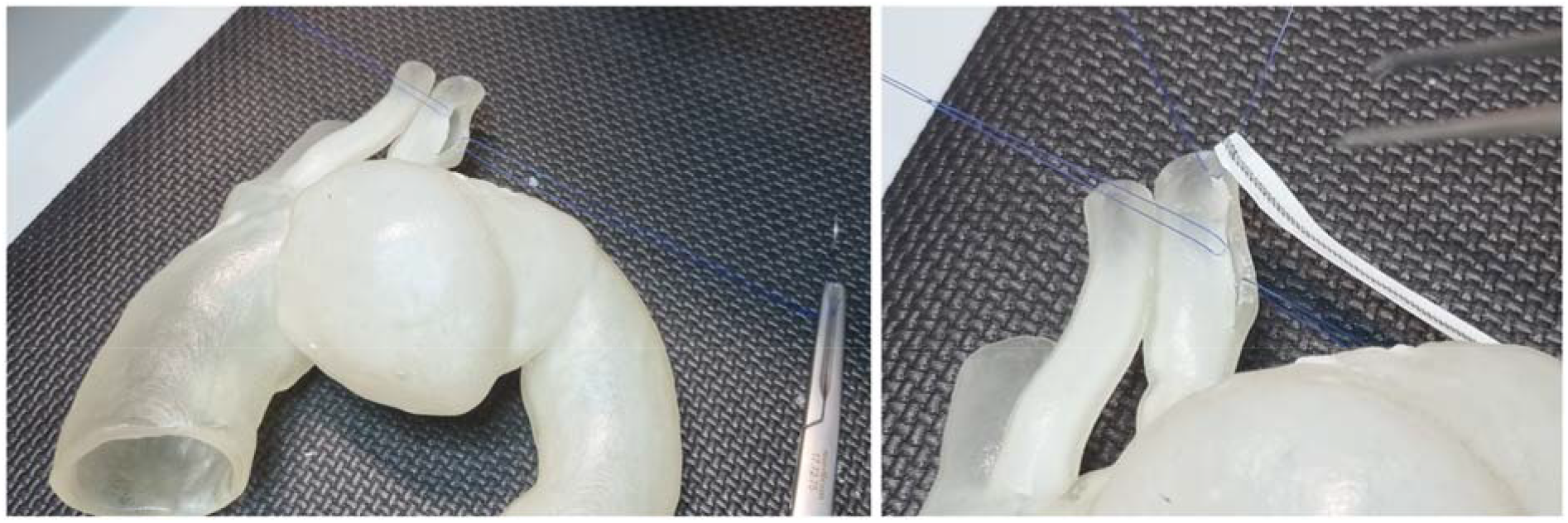
Two steps of the simulated procedure on C: cut and opening of the segments of the left subclavian artery (left) and placement of a synthetic patch (right). A tearing around stitch between the patch and the 3DP material is visible.

The morphology and the feeling of the model was considered good. Both A and C was properly cleaned from external Heavy Support and internal GelSupport without major issues. A small delamination at the level of the junction between the aneurismatic sac and the aortic arch was present in C model, due to the strong bending of the tissue at that level.

The simulation of the real procedure raised additional issues.

Performing the realistic procedure on the surgical approach on the aneurism, we observed several failures of the stitches. This is due to the fact that multiple stitches are made before tightening. When the stitch is finally pulled, then a high friction between the filament and the tissue is generated, causing the failure of the stitch. On the left subclavian artery, the simulation included the cut of the vessel and the placement of two stitches on the segments in order to keep the vessel opened attaching a clamp as a weight (Figure 13): the stitch withstood for a little the heavy of the clamp and failed. The most critical aspect of this procedure is the first stitch made to suture the patch. As visible in Figure 13 it is made along the line of the cut. This cause immediate failure due to delamination since this is the most critical situation for the material to withstand the stitch.

On C, the prolonged clamping of the vessels also resulted in a failure of the material. According to surgeons, A was considered representative of a calcific aorta, while C was evaluated as more in line with a normal tissue.

### 4.2. Abdominal aorta aneurism open surgery

A severe abdominal aorta aneurism was selected for the simulation. The aneurismatic sac reached a maximum diameter of 84 mm. The model was used just to test a simplified reconstruction of the right renal artery with a Gore-Tex Hybrid prosthesis. The anatomical model is made with a vessel wall of variable thickness, as shown in Figure 14. The model was printed using C for thoracic aorta (for the 2 mm thick tract above celiac trunk) and A from the celiac trunk on. The model was successfully printed with GelSupport in the inner lumen, despite the extremely high lumen diameter. The stitches were made with a Prolene 4/0 surgical thread. The procedure was carried out by two vascular surgeons.

**Figure 14:**
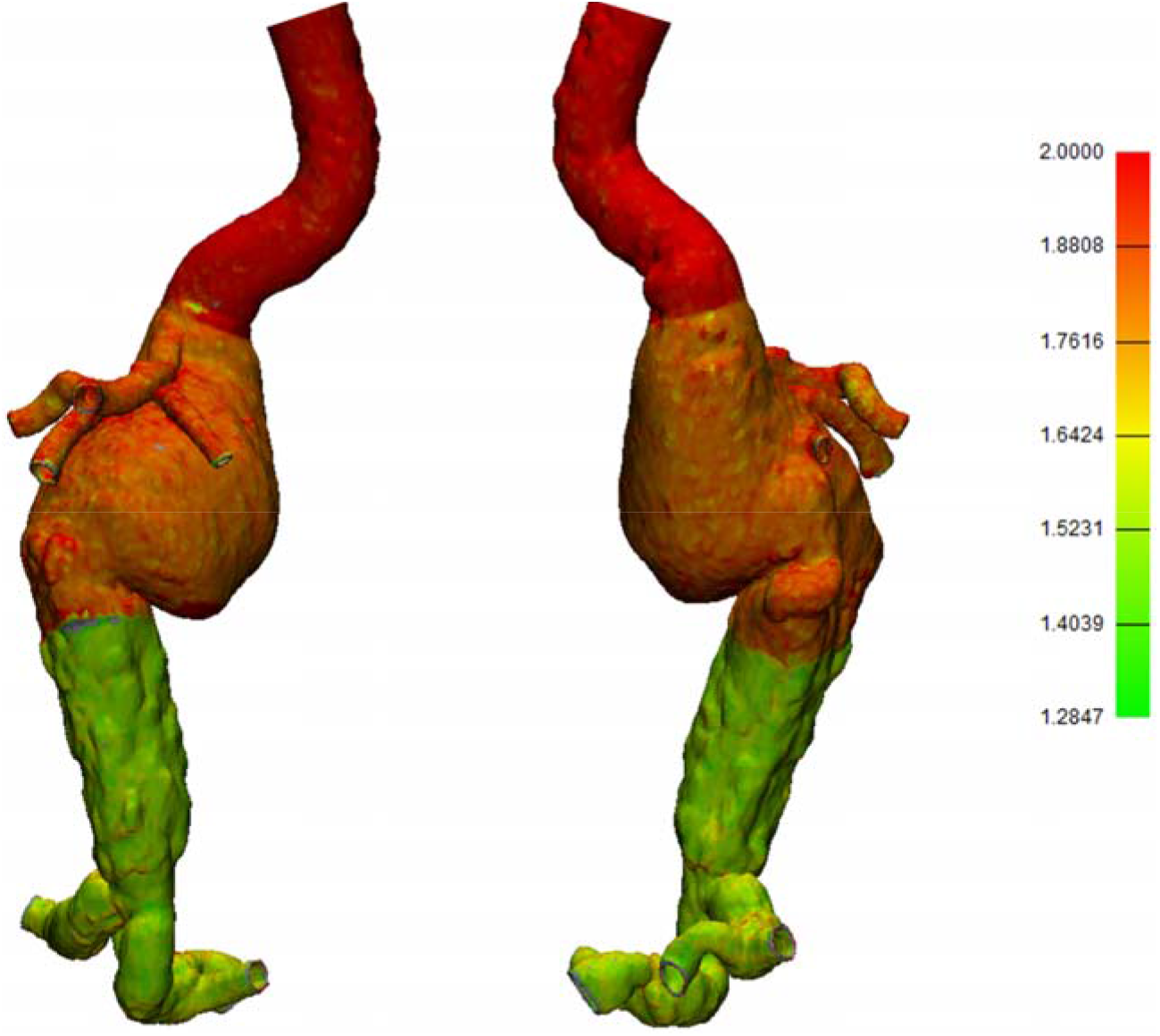
Abdominal aneurism model wall thicknesses. The model was printed using C for the thoracic aorta (red segment of 2 mm of wall thickness) while the remaining part was printed with A.

**Figure 14:**
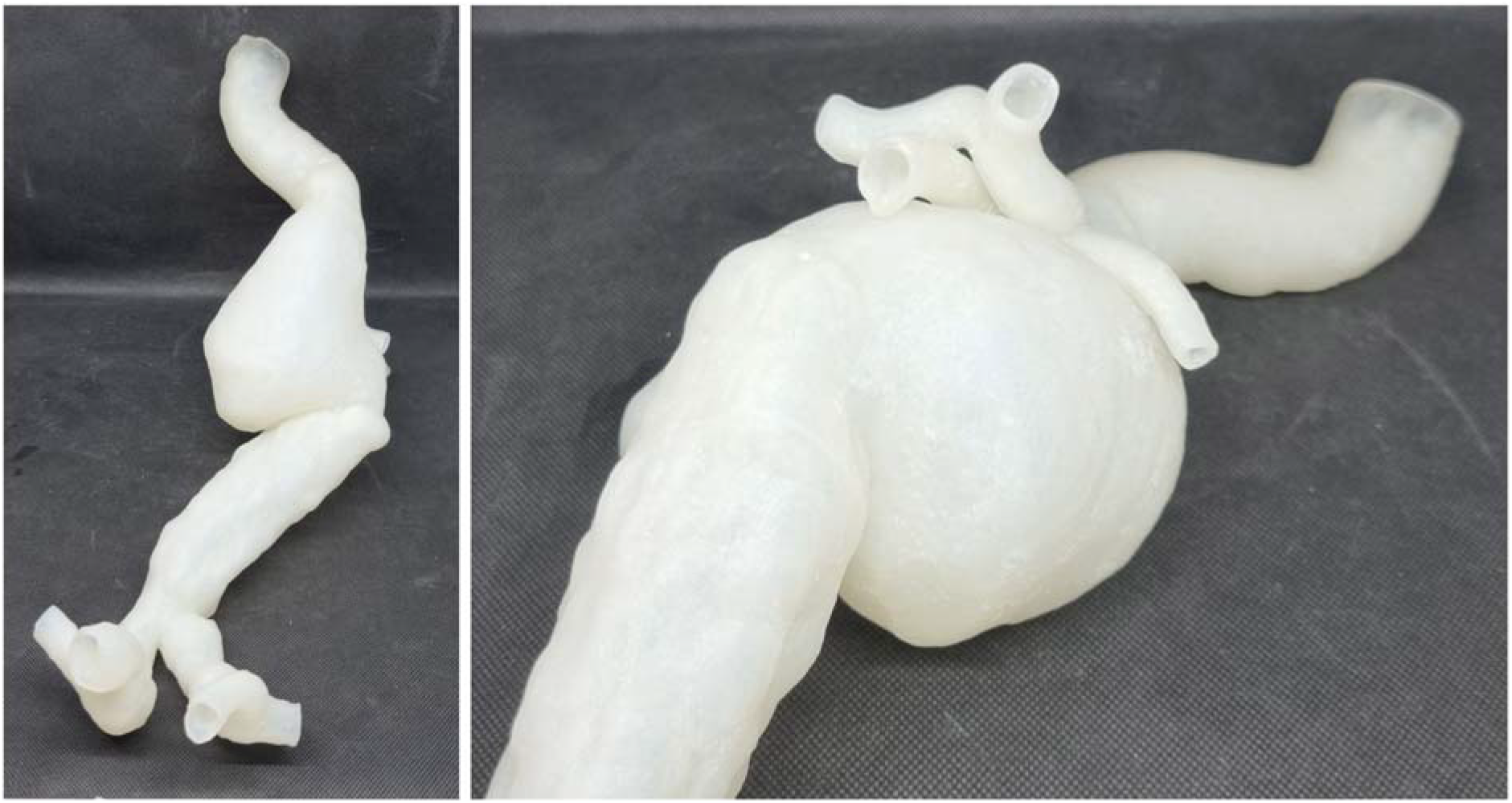
Abdominal aneurism model printed with A and C.

The morphology and the feeling of the model was considered good. Both A and C segments of the model were properly cleaned from external Heavy Support and internal GelSupport without any issue. The main interest on this model was to test the feasibility of the printing and cleaning. The overall cleaning was completed in about 3.5 hours. A small rupture (about 2 mm) at the level of the junction between the aneurismatic sac and the abdominal aorta was present, due to the strong bending of the tissue at that level. The procedure includes a circular stitch between the prosthesis and the graft. Due to the type of suture, here the A performed better in withstanding the stitch.

**Figure 16:**
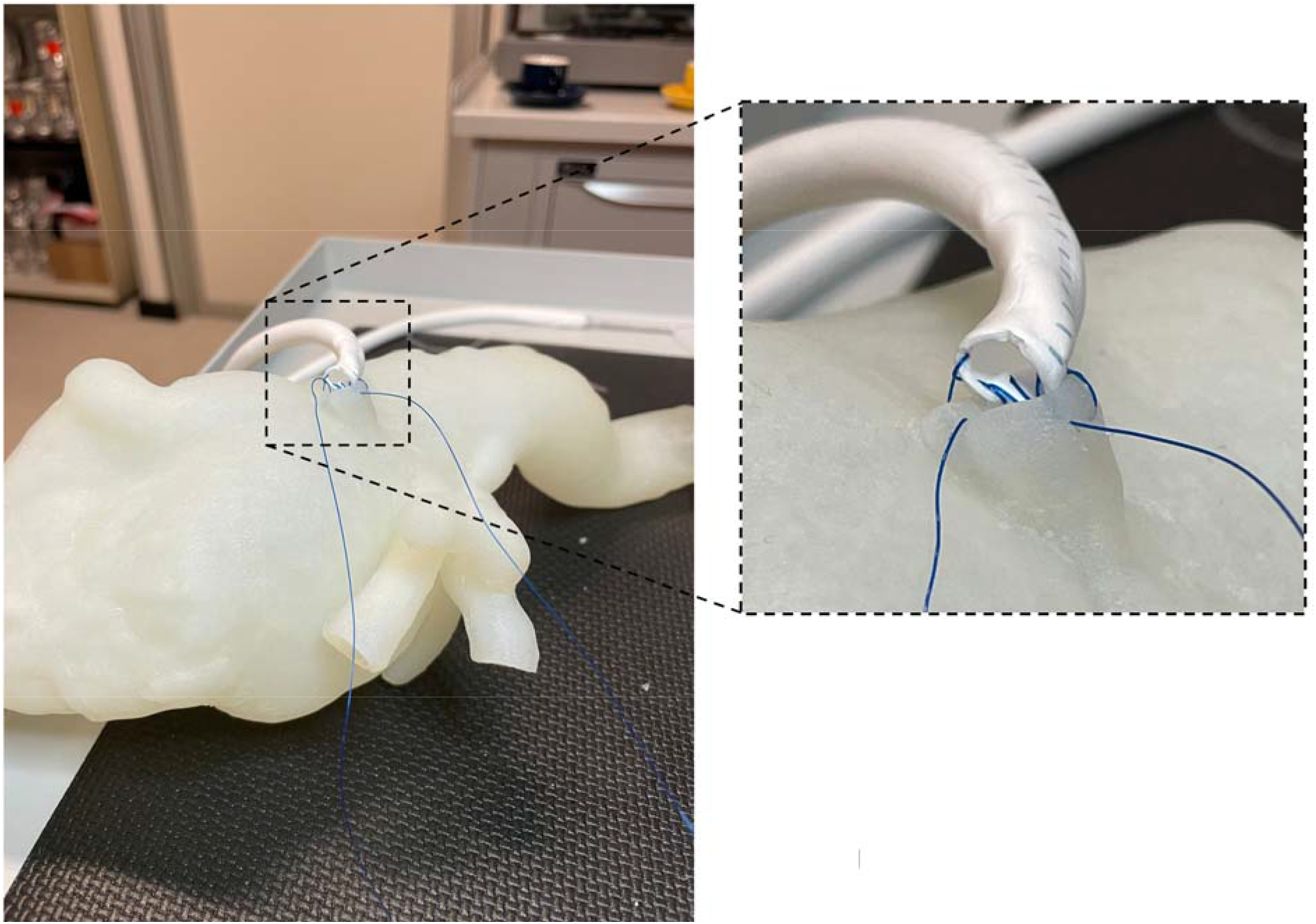
Circular stitch between the right renal artery and the Gore-tex graft.

### 4.3. Kidney transplantation in simulated environment

This surgery involves the anastomosis between donor renal arteries and veins and receivers’ iliac vessels. A complete 3DP simulation platform for kidney transplantation procedure has been used in this case [3]; all the involved anatomical structures were reconstructed from CT images of average adult patients, in order to perfectly reproduce the real morphology of the pelvic area (Figure 17).

**Figure 17:**
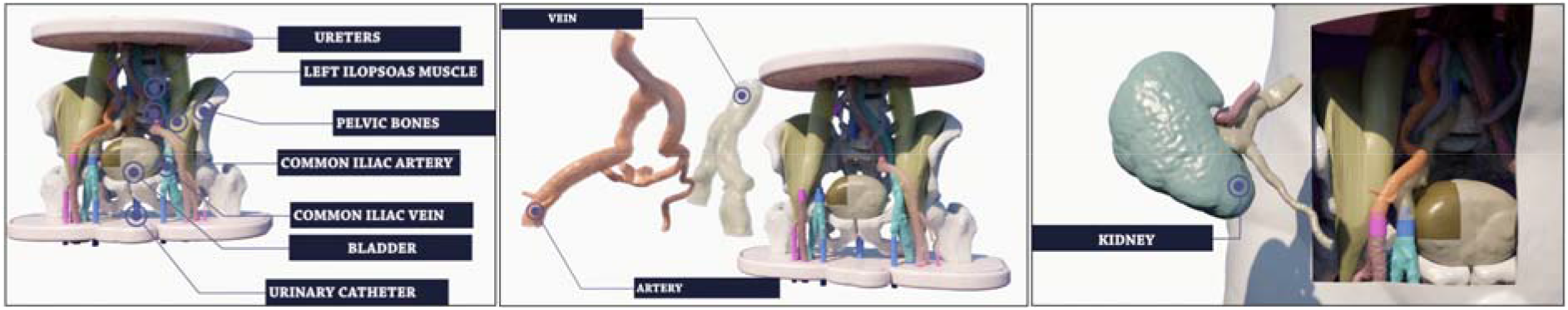
3D Virtual Rendering of the kidney simulator and a detail of receiving removable structures (iliac artery and vein) and donor removable structure (kidney).

The simulator is composed of fixed and removable parts; fixed structures – namely cutaneous surface, bones, ileo-psoas muscles, bladder, ureters and the left part of arterial and venous systems – were printed with suitable materials using a Objet260 Connex3 (Stratasys^®^).

Two sets of removable parts – which include both the receiving right iliac vessels and the donor renal artery and vein – are all printed entirely with A and C materials respectively. Receiving vessels are attached via sealing tube-connectors to the respective fixed proximal and distal vascular segments of the simulator, while donor vessels are connected to the kidney parenchyma in order to faithfully reproduce the transplantation procedure.

An expert surgeon performed the receiving-donor vessels’ anastomosis using a non-absorbable polypropylene/polyethylene 5/0 monofilament with a 3/8 18mm cylindrical needle.

The morphology and the feeling of the models was considered good. Both A and C sets of models were properly cleaned from support with all the available support strategies. It is anyway not recommended to use Standard/Heavy Support for the inner lumen support in very subtle vessels (0.7 −1 mm) with caliper > 5 mm, especially if they have a long extension as in the iliac vessels here used for the simulation (Figure 18). It should be noted that the same set of anatomical models was previously produced for the tests described in [3] using a combination of Agilus30 and VeroMagenta/VeroCyan. In that case, several delaminations of the models occurred during the cleaning. Vessels had to be covered with latex in order to perform the procedure.

**Figure 18:**
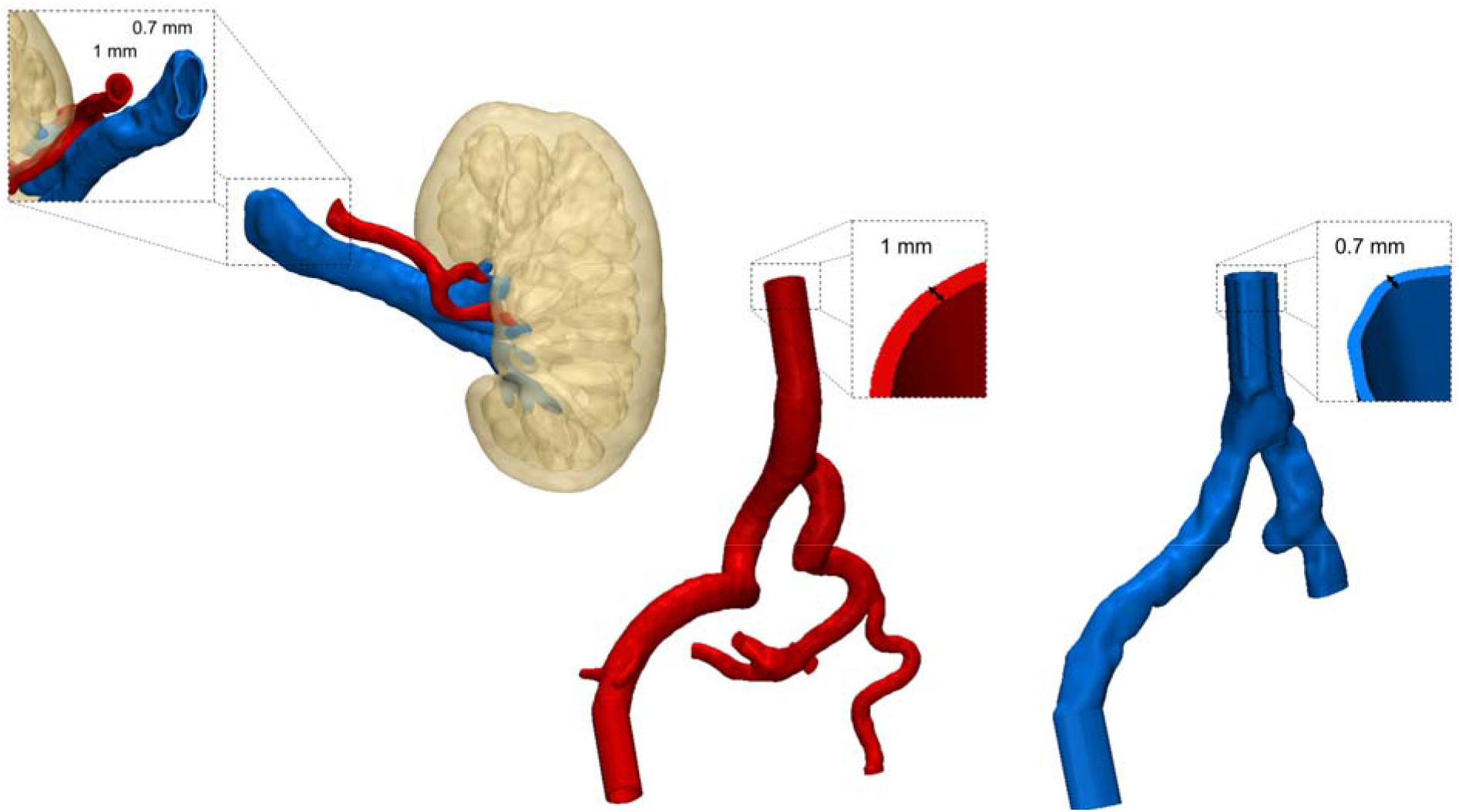
Wall thickness of receiver’s (iliac artery and vein) and donor’s (renal artery and vein) vessels.

During the testing of both A and C models we observed several failures of the stitches. As in the left subclavian artery, the most critical aspect of this procedure is the first stitch made to suture the renal artery to the iliac artery. As visible in Figure 19, the stitch is made along the line of the cut of the iliac artery. This cause immediate failure due to delamination since this is the most critical situation for the material to withstand the stitch. Moreover, the cut itself tends to extend over the original length due to the tearing of the material (Figure 19 - right). On the contrary, circular stitches perform better, especially after the first stitches are made, since the load start to be distributed among more stitches.

**Figure 19:**
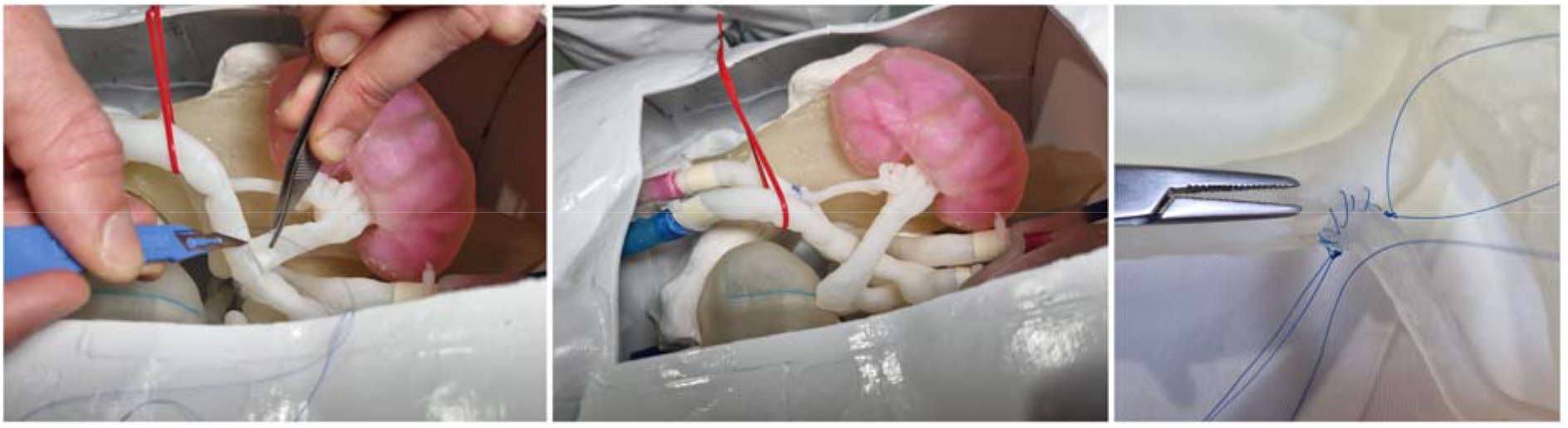
Details of the simulated procedure on A: cut of the iliac artery and suture of the renal artery.

## 4. Conclusions

The results of the mechanical and production tests identified in the A and C formulations the most promising solutions to be used for anastomoses simulation. Clinical applicative tests, specifically selected to challenge the new materials, raised additional issues on the performance of the materials to be considered for future developments.

The developed framework enabled us to deeply investigate and characterize the behavior of 3DP materials during suturing. Although additional work should be carried out to optimize their withstanding of the real applicative simulation, important advances have been carried out.

- A and C can be printed with inner GelSupport even in extremely critical situation (up to 84 mm of inner lumen diameter and 1.8 mm of wall thickness) enabling the complete cleaning of the inner lumen. This aspect is fundamental to pave the way to complex cases simulations. Accordingly, the use of GelSupport is highly recommended when producing anatomical models with such materials. External Heavy Support structure is suggested, especially for tortuous vessels extending a lot through layers, since it was safely cleaned.
- Clinical applicative cases highlighted how issues are still present in terms of failure of the stitches. This reflects the still unmet replica of the elongation of biological tissue and it deserves further improvement to come up whit a realistic simulation.

In view of all the gathered results, we can identify the following destination of use of the 2 developed materials.

- **A** can be selected to simulate more subtle vessels, as abdominal aorta or minor branches. This material exhibits higher unit peak force in puncture tests; thus it should be employed in lower thicknesses (about 1 mm) which are proper features of minor vessels.
- **C** can be selected to mimic medium strength vessels characterized by a higher thickness, as aortic arch and thoracic aorta. Regardless to the higher thickness, the material can give a good feeling in terms of perceived flexibility and softness.

## Acknowledgments

The authors want to thank the research group members who supported the testing campaign. Dr. Gianluca Alaimo and Martina Golosio, for mechanical testing, Virginia Gallo, MD, Tim Mandigers, MD, Sara Allievi, MD, for suture tests.

## Bibliography

[1] Lu J, Cuff RF, Mansour MA. Simulation in surgical education. Am J Surg. 2021;221(3):509–514.

[2] Araujo SEA, Perez RO, Klajner S. Role of Simulation-Based Training in Minimally Invasive and Robotic Colorectal Surgery. Clin Colon Rectal Surg. 2021;34(3):136–143.

[3] Peri A, Marconi S, Gallo V, Mauri V, Negrello E, Abelli M, Ticozzelli E, Caserini O, Pugliese L, Auricchio F, Pietrabissa A. Three-D-printed simulator for kidney transplantation. Surg Endosc. 2021 Nov 15. doi: 10.1007/s00464-021-08788-1. Epub ahead of print. PMID: 34782966.

[4] Ganguli A, Pagan-Diaz GJ, Grant L, Cvetkovic C, Bramlet M, Vozenilek J, Kesavadas T, Bashir R. 3D printing for preoperative planning and surgical training: a review. Biomed Microdevices. 2018 4;20(3):65.

[5] Pietrabissa A, Marconi S, Negrello E, Mauri V, Peri A, Pugliese L, Marone EM, Auricchio F. An overview on 3D printing for abdominal surgery. Surg Endosc. 2020 Jan;34(1):1–13. doi: 10.1007/s00464-019-07155-5. Epub 2019 Oct 11. PMID: 31605218; PMCID: PMC7175828.

[6] Bao, X. et al. (2016). “Experiment study on puncture force between MIS suture needle and soft tissue”.

[7] Maurin, B. et al. (2004). “In vivo study of forces during needle insertions”. Perspective in Image-Guided Surgery, pp. 415–422.

[8] Okamura, A.M. et al. (2004). “Force modeling for needle insertion into soft tissue”. IEEE transactions on biomedical engineering 51.10, pp. 1707–1716.

[9] Patel, H.B. et al. (2004). “Puncture resistance and stiffness of nitrile and latex dental examination gloves”. British dental journal 196.11, pp. 695–700.

[10] Nguyen, C.T. et al. (2009). “Puncture of elastomer membranes by medical needles. Part I: Mechanisms”. International journal of fracture 155.1, pp. 75–81.

[11] Podder, T. et al. (2006). “In vivo motion and force measurement of surgical needle intervention during prostate brachytherapy”. Medical physics 33.8, pp. 2915–2922.

[12] Pavicic, T. et al. (2019). “Arterial wall penetration forces in needles versus cannulas”. Plastic and reconstructive surgery 143.3, pp. 504–512.

[13] Zhang, G. et al. (2017). “Development of a penetration friction apparatus (PFA) to measure the frictional performance of surgical suture”. Journal of the mechanical behavior of biomedical materials 74, pp. 392–399.

[14] Zhai, J. et al. (2013). “A sensor for needle puncture force measurement during interventional radiological procedures”. Medical engineering & physics 35.3, pp. 350–356.

[15] Urrea, F. A et al. (2016). “Evaluation of the friction coefficient, the radial stress, and the damage work during needle insertions into agarose gels”. Journal of the Mechanical Behavior of Biomedical Materials 56, pp. 98–105.

[16] Jong, T.L. de et al. (2017). “PVA matches human liver in needle-tissue interaction”. Journal of the mechanical behavior of biomedical materials 69, pp. 223–228.

[17] Tan, Z. et al. (2018). “Composite hydrogel: A high fidelity soft tissue mimic for surgery”. Materials & Design 160, pp. 886–894.

[18] Casanova, F. et al. (2014). “In vivo evaluation of needle force and friction stress during insertion at varying insertion speed into the brain”. Journal of neuroscience methods 237, pp. 79–89.

[19] Westwood, J.D et al. (2005). “In vivo force during arterial interventional radiology needle puncture procedures”. Medicine Meets Virtual Reality: The Magical Next Becomes the Medical Now. Vol. 111, pp. 178–184.

[20] (2020). url: https://www.astm.org/Standards/F1342.htm (visited on 14/07/2020).

